# Comparative analysis of the chloroplast genome of *Paeonia suffruticosa* ‘Cun Song Ying’ reveals plastid characteristics and phylogenetic relationships

**DOI:** 10.1101/2025.10.10.681658

**Authors:** Wenzhen Cheng, Conghao Hong, Mingyu Li

**Affiliations:** College of Agriculture and Bioengineering, Heze University, Heze, 274000, China; Shanghai Construction Management Vocational and Technical College,Shanghai, 200232, China

**Author notes:** These authors contributed equally to this work (Wenzhen Cheng, Conghao Hong).

**Keywords:** *Paeonia*, chloroplast genome, phylogeny, codon usage

## Abstract

Chloroplast genomes provide valuable insights into plant evolution, phylogeny, and molecular breeding. Here, we present the first complete chloroplast genome of *Paeonia suffruticosa* ‘Cun Song Ying’. Using high-throughput sequencing combined with multiple bioinformatics approaches, we comprehensively characterized its chloroplast genome and compared it with those of closely related taxa. The chloroplast genome of ‘Cun Song Ying’ is 152,704 bp in length and exhibits the typical quadripartite structure, comprising a large single-copy region (LSC), a small single-copy region (SSC), and two inverted repeat regions (IRa and IRb). A total of 138 genes were annotated, including 4 pseudogenes and 134 functional genes, which consist of 87 protein-coding genes, 39 tRNAs, and 8 rRNAs. Phylogenetic analysis revealed that ‘Cun Song Ying’ shares the closest relationship with *P. suffruticosa* ‘Luo Yang Hong’. These findings provide new insights into the evolutionary history of peony plastids, and the identified highly variable regions may serve as potential DNA barcodes for cultivar identification.

## 1. Introduction

Chloroplasts are key organelles in plant cells responsible for photosynthesis and various metabolic pathways. Their origin can be traced back to an ancient endosymbiotic event with cyanobacteria approximately 1.5 billion years ago [1, 2]. During this evolutionary process, primitive cyanobacteria established a stable endosymbiotic relationship with host eukaryotic cells. Although their genomes underwent substantial reduction, resulting in only 110-130 genes remaining in the chloroplast genomes of most terrestrial plants [3], they retained their own replication and transcription systems and their genomic DNA was evolved into a distinct circular quadripartite structure [4]. This semi-autonomous nature makes chloroplast genomes valuable molecular markers for phylogenetic reconstruction, species identification, and crop genetic improvement, particularly in distinguishing closely related taxa and detecting hybridization events [5].

Due to their high copy number, maternal inheritance, and conserved gene content and organization, chloroplast genomes are powerful tools for studying phylogenetic relationships among closely related species [6-8]. In angiosperms, the typical chloroplast genome displays a highly conserved quadripartite structure in which a pair of 26 kb inverted repeats (IRs) divide the circular DNA molecule into large and small single-copy regions (LSC and SSC). This structure is important in maintaining genome stability and facilitating homologous recombination [9]. Recent studies have revealed structural variations across plant lineages, including IR boundary expansion or contraction, gene loss, and sequence inversion, which provide new perspectives for investigating plant adaptive evolution [10-12].

Many species in the *Paeoniaceae* family hold not only exceptional ornamental value but also significant medicinal importance. Among them, *Paeonia suffruticosa* ‘Cun Song Ying’ is recognized as an important cultivated variety owing to its distinctive pink flowers and robust growth performance. Although advances in high-throughput sequencing have led to the inclusion of more than 7,000 plant chloroplast genomes in the NCBI Organelle Genome Database [13], the chloroplast genome features and phylogenetic placement of this cultivar remain unexplored. In this study, we report for the first time the complete chloroplast genome sequence of ‘Cun Song Ying’ (152,704 bp). Its genome exhibits the canonical quadripartite structure, comprising an LSC region (84,365 bp), an SSC region (17,047 bp), and two IR regions (25,646 bp each), and encodes 134 functional genes, including 87 protein-coding genes, 39 tRNA genes, and 8 rRNA genes. Comparative genomic analyses clarify structural variation patterns within *Paeoniaceae* chloroplast genomes, provide molecular evidence for phylogenetic inference, and establish a foundation for future molecular breeding efforts such as flower color improvement.

Although the genetic code is highly conserved, the frequency of synonymous codon usage varies substantially across species, a phenomenon known as codon usage bias [14]. Population genetics studies indicate that although synonymous sites are under relatively weak selective pressure, codon usage patterns reflect the combined effects of mutation, natural selection, and genetic drift over long-term evolution [15]. Codon usage bias, defined as the non-random use of synonymous codons encoding the same amino acid, is shaped by complex evolutionary mechanisms. Investigating codon usage bias in plant chloroplast genomes can provide insights into phylogenetic relationships, horizontal gene transfer, and molecular evolution, while also contributing to the optimization of gene expression, enhancement of genetic transformation, and theoretical guidance for species conservation [16,17].

In this study, we assembled, annotated, and performed comparative analyses of the chloroplast genome of *Paeonia suffruticosa* ‘Cun Song Ying’. We systematically examined its structural features, distribution of repetitive sequences, codon usage bias, and IR boundary variation. In addition, we constructed a phylogenetic tree including closely related species to clarify the evolutionary characteristics of its plastid genome and its phylogenetic position within *Paeoniaceae*. Our findings provide molecular markers for species identification and genetic diversity analysis of ‘Cun Song Ying’, contribute new genomic evidence for taxonomic revision and evolutionary history reconstruction in *Paeoniaceae*, and establish a theoretical foundation for the conservation and future utilization of this cultivar.

## 2 Materials and Methods

### 2.1 Sampling, DNA Extraction, and Chloroplast Genome Sequencing

Fresh leaf samples of *Paeonia suffruticosa* ‘Cun Song Ying’ were collected at Heze University. The samples were immediately frozen in liquid nitrogen and stored at –80 °C. Total genomic DNA was extracted using the HiPure SF Plant DNA Mini Kit (Magen). After DNA end repair and 3’-end adenylation, sequencing adapters were ligated, and target fragments were recovered using magnetic bead purification. The fragments were amplified by PCR to construct sequencing libraries. Library quality was assessed prior to sequencing, and qualified libraries were sequenced on the Illumina NovaSeq X Plus platform.

### 2.2 Sequencing Data Quality Control, Assembly, and Chloroplast Gene Annotation

To ensure the accuracy of downstream analyses, raw reads were filtered using Cutadapt (v1.16) [18] to remove adapter-containing reads, reads with more than 10% ambiguous bases (N), and reads in which low-quality bases (Phred score < 5) accounted for more than 50% of the sequence length. After stringent filtering, clean reads were retained for subsequent analyses. Base composition distribution was examined to assess possible AT/GC bias. Quality profiles of sequencing reads across all cycles were evaluated using FastQC (v0.11.4) (http://www.bioinformatics.babraham.ac.uk/projects/fastqc/), providing an overview of sequencing data quality.

Genome assembly was performed using NOVOPlasty (v4.2) (https://github.com/ndierckx/novoplasty) [19], Fast-Plast (v1.2.8) (https://github.com/mrmckain/Fast-Plast), and GetOrganelle (v1.7.0+) (https://github.com/Kinggerm/GetOrganelle) [20]. The assembly with the best overall performance was selected as the final result. Annotation of the assembled chloroplast genome was conducted using PGA [21] and GeSeq(https://chlorobox.mpimp-golm.mpg.de/geseq.html) [22], followed by manual correction of all annotation results.

Predicted protein-coding sequences were compared against protein databases to obtain functional annotations. To ensure biological relevance, only the best alignment result for each gene was retained. Functional annotation databases included NR (http://www.ncbi.nlm.nih.gov/), Swiss-Prot (http://www.ebi.ac.uk/uniprot), eggNOG (http://eggnogdb.embl.de/), KEGG (http://www.genome.jp/kegg/), and GO (http://geneontology.org/). A circular gene map of the chloroplast genome was generated using the online tool OGDRAW [23] (https://chlorobox.mpimp-golm.mpg.de/OGDraw.html).

### 2.3 Comparative Genomic Analysis

The chloroplast genome of *Paeonia suffruticosa* ‘Cun Song Ying’ was compared with 15 published *Paeoniaceae* chloroplast genomes (Table S1). Sequence similarity was assessed using mVISTA in LAGAN mode, which enables accurate multiple alignment regardless of potential inversions [24]. Genome sequences were aligned using MAFFT (7.525) [25], and nucleotide diversity (π) between ‘Cun Song Ying’ and other *Paeoniaceae* species was estimated with DnaSP v6.12 using a sliding window analysis (window length: 600 bp; step size: 200 bp) [26]. Results were visualized using ggplot2 package in R (R-4.2.3).

To detect potential genome rearrangements, complete chloroplast genome alignments were conducted with ProgressiveMauve v.1.1.3 implemented in Geneious Prime 2025.2.1 [27]. IRplus (https://irscope.shinyapps.io/IRplus/) was used to examine the contraction or expansion of inverted repeat boundaries [28].

### 2.4 Analysis of Repetitive Sequences

Repetitive sequences, including direct, reverse, palindromic, and complementary repeats, were identified using REPuter (https://bibiserv.cebitec.uni-bielefeld.de/reputer) [29] with the following parameters: minimum repeat size of 30 bp, maximum repeat length of 5,000 bp, Hamming distance of 3,000 bp, and sequence identity ≥ 90%. Simple sequence repeats (SSRs) were detected using MISA-web (http://pgrc.ipk-gatersleben.de/misa/), with the following minimum thresholds: 10 repeat units for mononucleotides, 5 for dinucleotides, 4 for trinucleotides, and 3 for tetranucleotides, pentanucleotides, and hexanucleotides. Relative synonymous codon usage (RSCU) values were calculated using CodonW (v1.4.4) [30].

### 2.5 Phylogenetic Analysis

Phylogenetic relationships were inferred using the shared protein-coding genes (PCGs) from 16 *Paeoniaceae* chloroplast genomes. *Paeonia sterniana* and *P. veitchii* were selected as outgroups (Table S1). Gene sequences were extracted, concatenated, and aligned with MAFFT (7.525) in ClustalW mode, followed by manual inspection to ensure reading frame accuracy. Maximum likelihood (ML) analyses were performed with RAxML v.8.2.4 using 1,000 bootstrap replicates to assess branch support [31]. Bayesian inference (BI) was conducted in MrBayes v.3.2.7 [32] with 1,000,000 Markov chain Monte Carlo (MCMC) generations. The first 25% of sampled trees were discarded as burn-in, and the remaining trees were used to generate a majority-rule consensus tree.

For both ML and BI analyses, the best-fitting nucleotide substitution model was determined using jModelTest v.2.1.9 [33] under the Akaike Information Criterion (AIC). Phylogenetic trees were visualized with FigTree v.1.4.5 (http://tree.bio.ed.ac.uk/software/figtree/).

## 3 Results

### 3.1 Assembly and annotation of the *Paeonia suffruticosa* ‘Cun Song Ying’ chloroplast genome

Illumina sequencing generated 8.5 Gb of raw data, with a Q30 value of 97.64%, indicating high-quality sequencing output. The assembled chloroplast genome of *Paeonia suffruticosa* ‘Cun Song Ying’ was 152,704 bp in length and exhibited the typical quadripartite structure. The LSC region (84,365 bp) and SSC region (17,047 bp) were separated by two IR regions (25,646 bp each) (Figure 1, Table S2).

**Figure 1.**
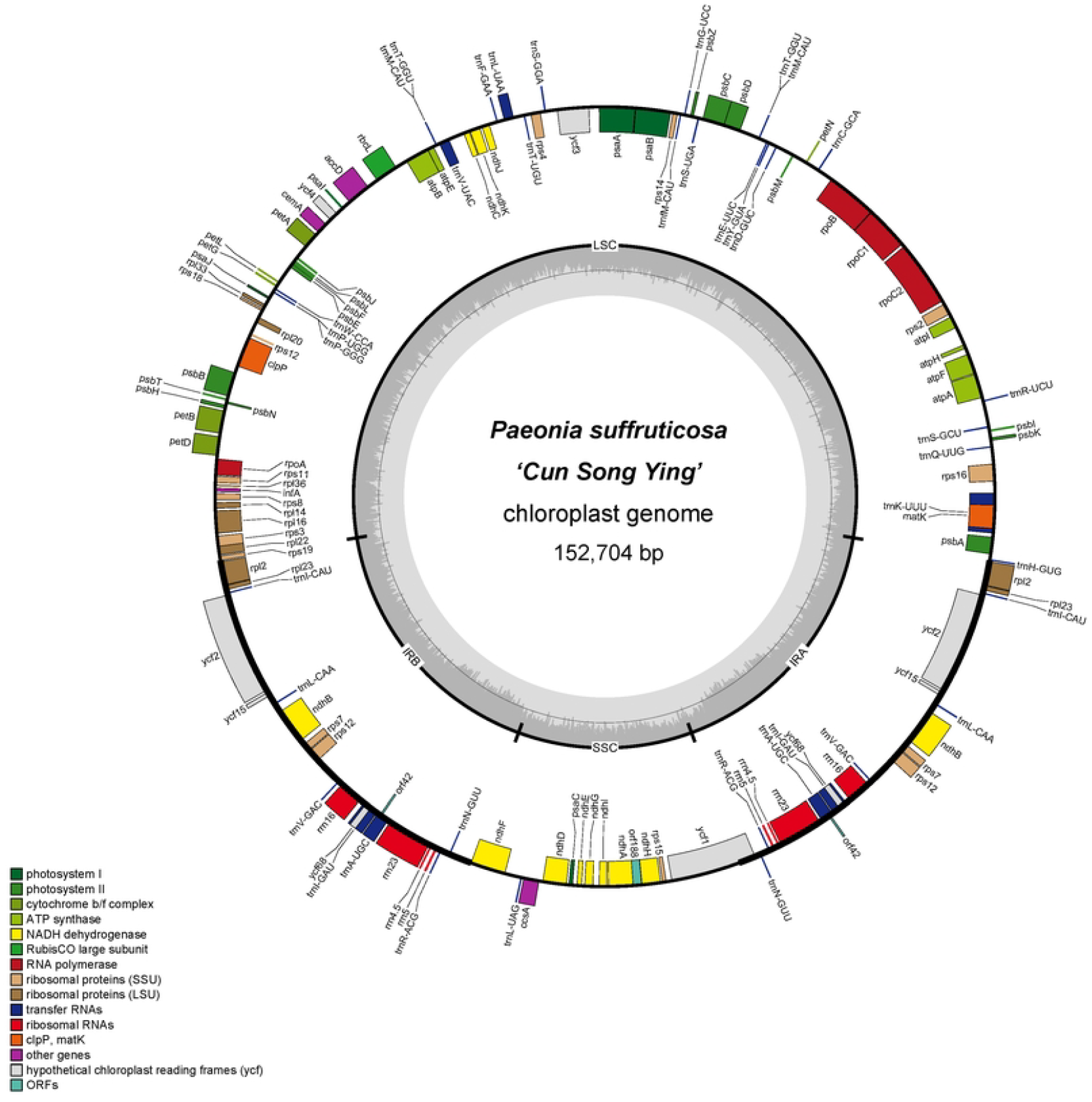
Chloroplast genome map of *Paeonia suffruticosa* ‘Cun Song Ying’. Genes inside the circle are transcribed clockwise, while those outside are transcribed counterclockwise. Functional groups are indicated by color. The inner dark gray circle represents GC content and the light gray circle represents AT content.

A total of 134 functional genes were annotated, including 87 protein-coding genes (PCGs), 39 tRNA genes, and 8 rRNA genes. Among these, 74 genes were associated with self-replication: 9 related to the large ribosomal subunit, 14 to the small ribosomal subunit, 4 encoding RNA polymerase subunits, 8 encoding rRNAs, and 39 encoding tRNAs. In addition, 45 genes were involved in photosynthesis, including 5 encoding subunits of photosystem I, 15 for photosystem II, 12 for NADH dehydrogenase, 6 for the cytochrome b/f complex, 1 for the large subunit of Rubisco, and 6 for ATP synthase.

Twelve genes were annotated with other or putative functions (*matK, clpP, cemA, accD, ccsA*) or unknown functions (*ycf1, ycf2, ycf3, ycf4, ycf15*). Twenty-two genes contained introns: 18 harbored a single intron (*atpF, ndhA, ndhB ×2, petB, petD, rpl16, rpl2 ×2, rpoC1, rps16, trnA-UGC ×2, trnI-GAU ×2, trnK-UUU, trnL-UAA, trnV-UAC*), while four genes (*clpP, rps12 ×2, ycf3*) contained two introns (Table S3).

### 3.2 Comparative analysis and nucleotide polymorphism

Sequence similarity across chloroplast genomes of *Paeoniaceae*, including ‘Cun Song Ying’, was assessed using mVISTA. Coding regions are more conserved than noncoding regions (Figure S1). No major structural rearrangements were detected across the compared genomes.

Nucleotide polymorphism (π) was calculated for 14 *Paeoniaceae* chloroplast genomes. The average π value was 0.0031, ranging from 0 to 0.01658 (Figure 2). IR regions were the most conserved (average π = 0.0106), whereas higher variability was observed in the LSC (average π = 0.0391) and SSC regions (average π = 0.0458).

**Figure 2.**
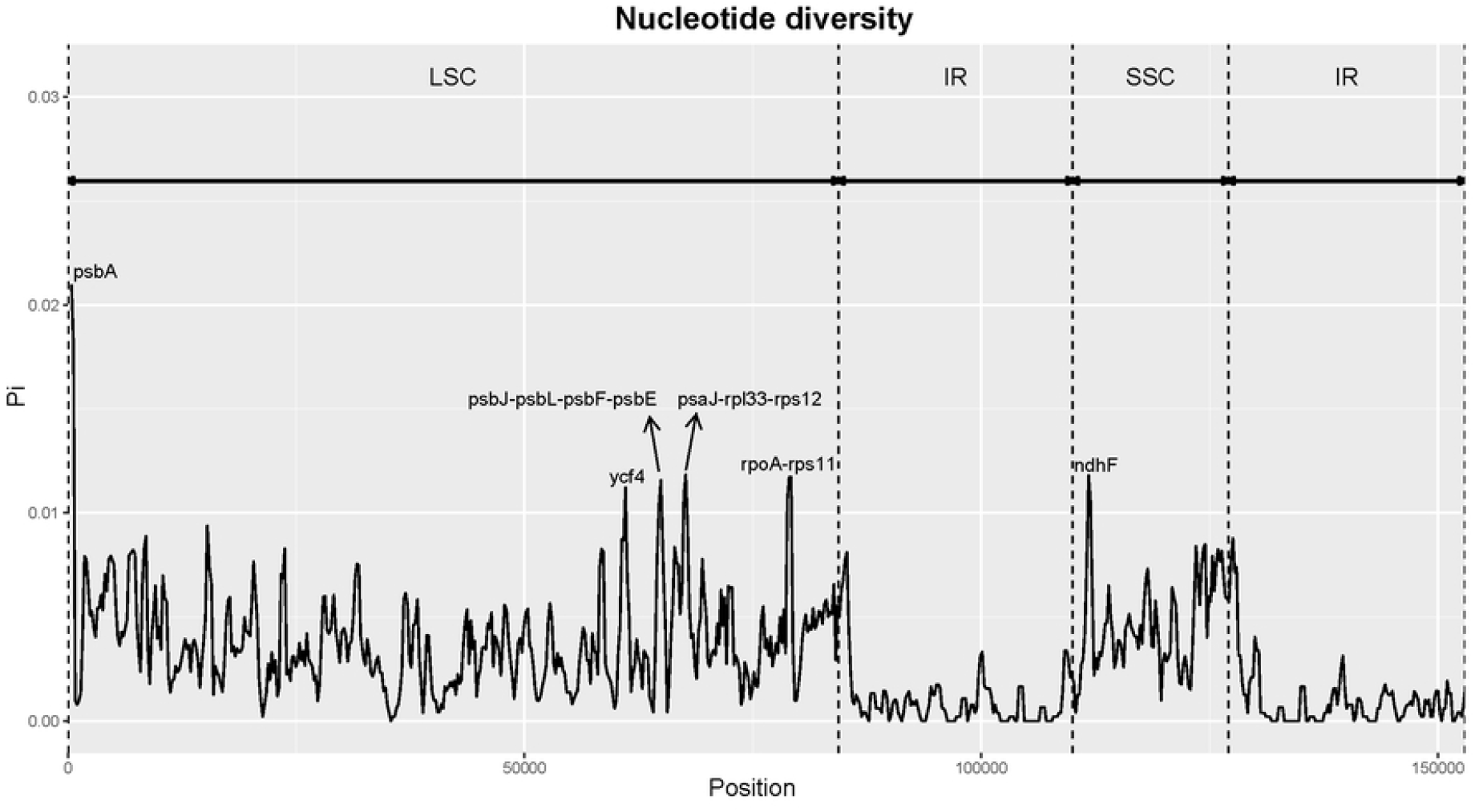
Nucleotide diversity (π) across 15 peonies chloroplast genomes.

Six hypervariable regions (π > 0.01) were identified: *psbA* (π = 0.02097), *ycf4* (π = 0.01122), *psbJ–psbL–psbF–psbE* (π = 0.01158), *psaJ–rpl33–rps12* (π = 0.01186), *rpoA– rps11* (π = 0.01174), and *ndhF* (π = 0.01181). These regions represent promising candidates for molecular markers in peony phylogenetics and population genetics(Figure 2).

### 3.3 Tandem repeats and simple sequence repeats (SSRs)

A total of 43 dispersed repeats were identified in *Paeonia suffruticosa* ‘Cun Song Ying’, including 21 palindromic repeats and 22 forward repeats. In contrast, 616 repeats were detected across the other 15 peonies, comprising 300 palindromic, 309 forward, and 7 reverse repeats (Figure 3A, Table S4). Repeat lengths ranged from 30 to 79 bp. The most frequent repeat sizes were 31, 32, 34, 35, 39, 52, and 54 bp, which were shared across all species. Repeats of 38 bp (present in 14 species) and 41 bp (present in 13 species) were also common (Figure 3B).

**Figure 3.**
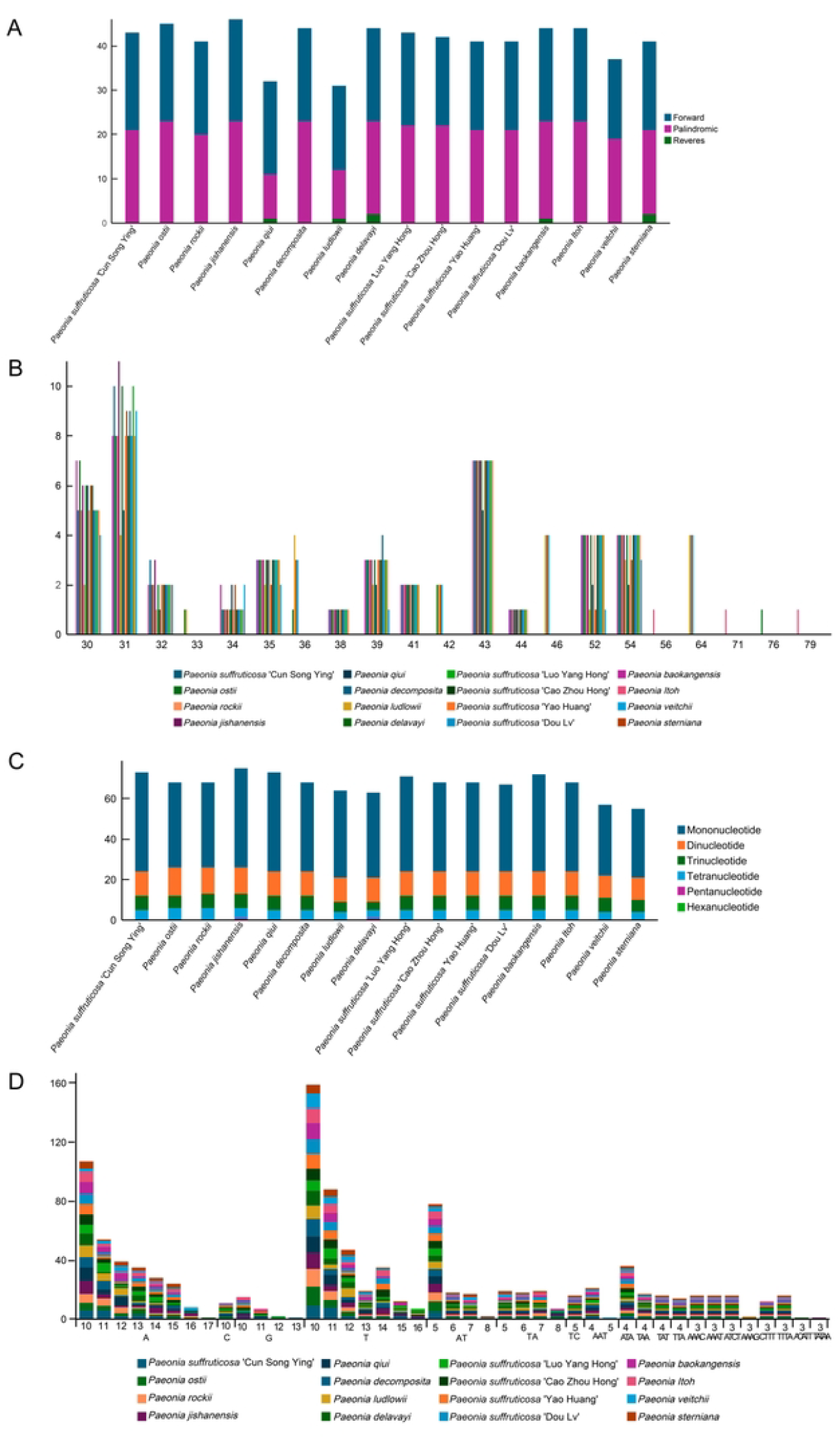
Repeat sequence analyses across 16 *Paeoniaceae* chloroplast genomes. (A) Distribution of tandem repeat types. (B) Frequency of tandem repeats of different lengths. (C) Distribution of SSR motifs. (D) Frequency of SSR motif types.

In *Paeonia suffruticosa* ‘Cun Song Ying’, 73 SSRs were identified, including 49 mononucleotide, 12 dinucleotide, 7 trinucleotide, and 5 tetranucleotide motifs (Figure 3C, TableS5). Among *Paeonia* species, the total number of SSRs ranged from 55 in *P. sterniana* to 75 in *P. jishanensis*. Species such as *P. sterniana* and *P. veitchii* contained relatively few SSRs (55–57). Pentanucleotide SSRs were observed only in *P. delavayi* and *P. jishanensis* (Figure 3D, TableS5).

Boundary analyses indicated that in ‘Cun Song Ying’, the LSC/IRb junction was located within *rps19*, the LSC/IRa junction was adjacent to *trnH*, the IRa/SSC junction intersected *ycf1*, and the IRb/SSC boundary was at *ndhF*. The IR regions ranged from 24,863 to 25,651 bp across *Paeonia* species, with conserved gene content in all taxa examined (Figure 4).

**Figure 4.**
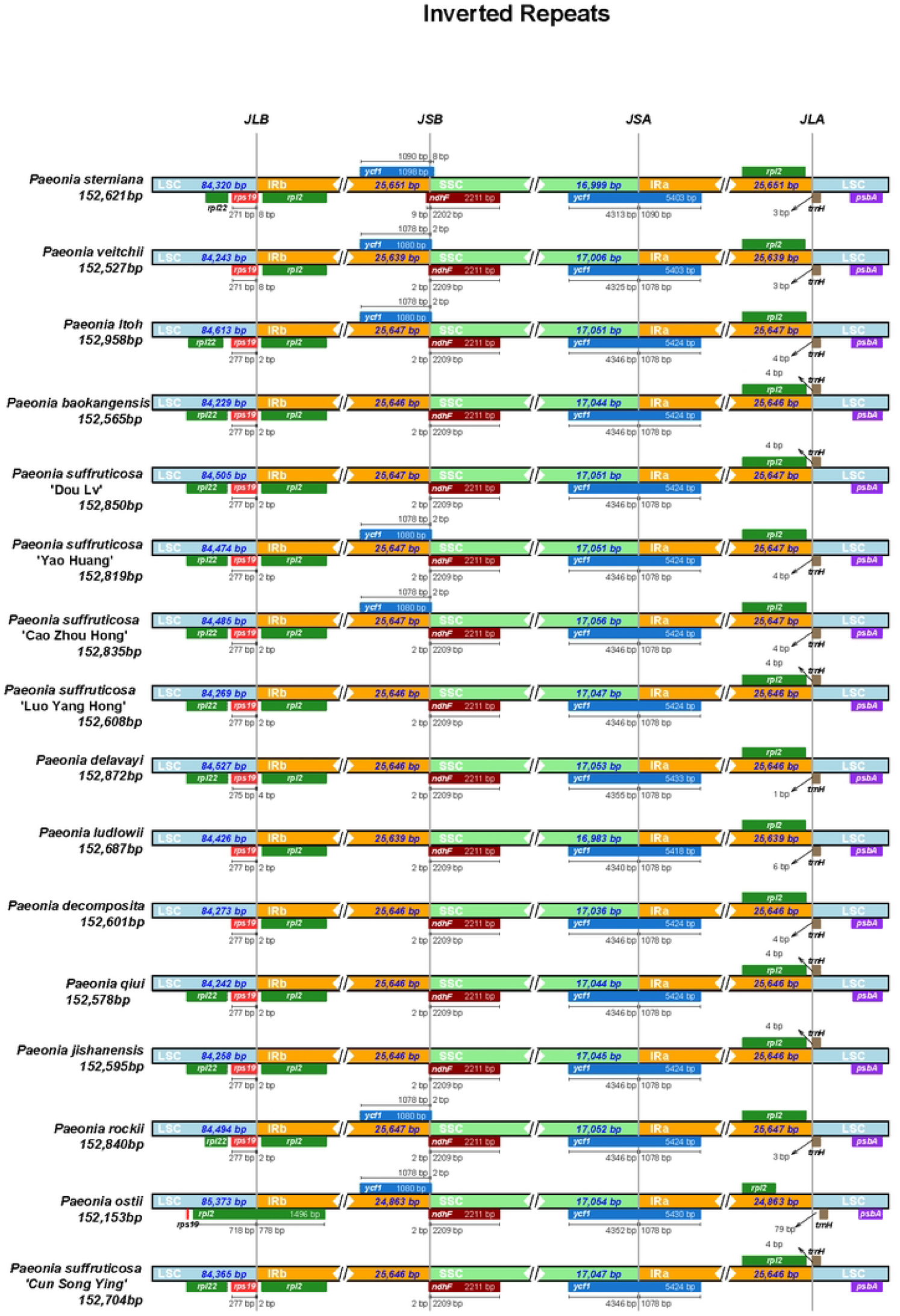
Comparison of LSC, IR, and SSC boundaries among *Paeoniaceae* species.

### 3.4 Relative synonymous codon usage (RSCU)

RSCU values were calculated from the complete coding sequence of the *Paeonia suffruticosa* ‘Cun Song Ying’ chloroplast genome, based on 69 PCGs and 24,297 codons. Leucine (Leu, 10.40%) was the most abundant amino acid, whereas cysteine (Cys, 1.13%) was the least frequent. The most frequently used codon was AUU (1,011 counts; encoding isoleucine), while UGC (71 counts; encoding cysteine) was the least common (Table S6).

Eighteen codons displayed codon usage bias with RSCU > 1.0. ATG and TGG showed no codon usage bias (RSCU = 1.0). Codon usage bias was particularly evident in leucine: UUA exhibited the highest RSCU value (1.97), while CUG had the lowest (0.37). Overall, codons ending in A/U were preferred over G/C-ending codons, consistent with trends reported for other *Paeonia* species (Figure 5).

**Figure 5.**
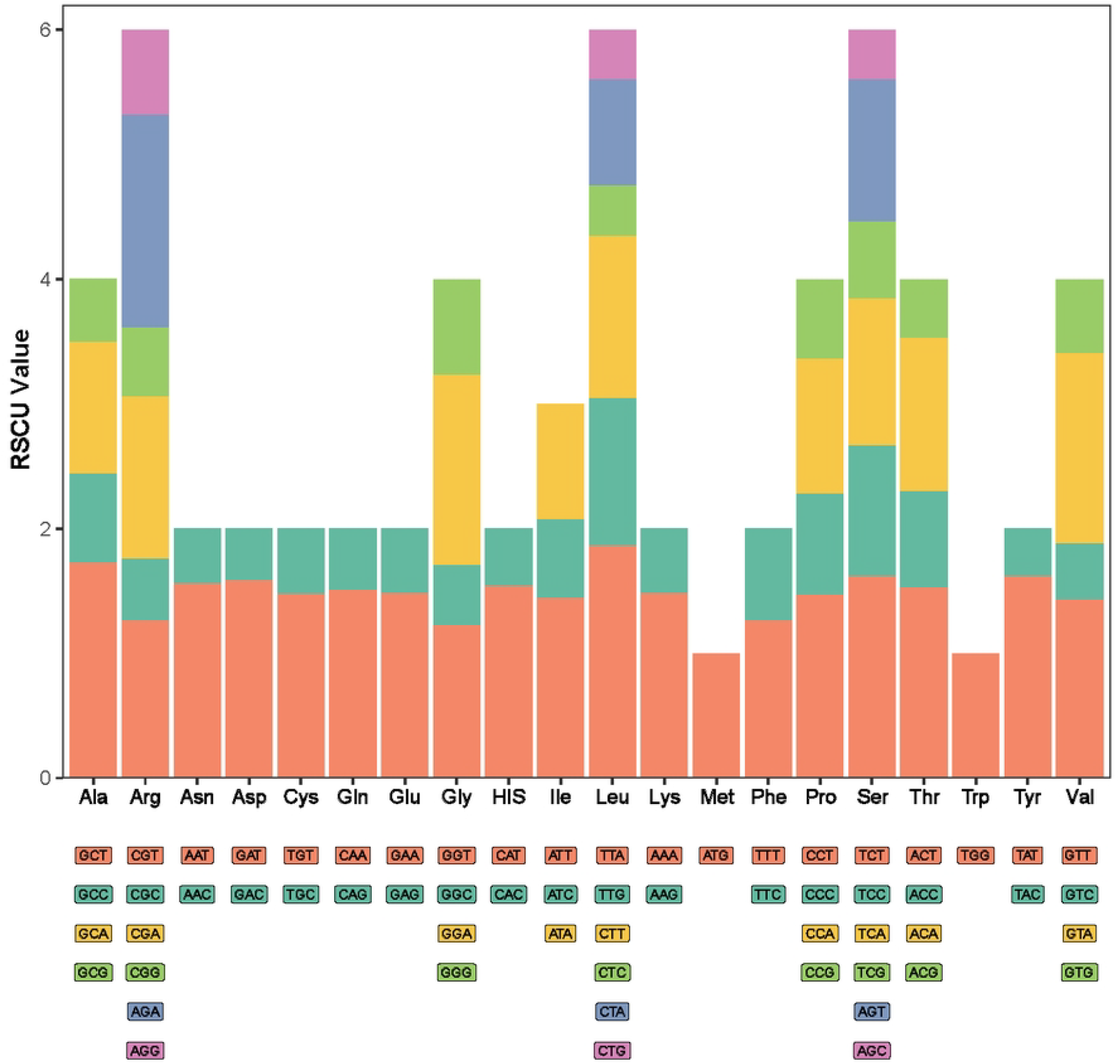
Relative synonymous codon usage (RSCU) values for 20 amino acids in chloroplast protein-coding genes of *Paeonia*. The x-axis represents amino acids, the y-axis represents RSCU values, and colors indicate different synonymous codons.

### 3.5 Phylogenetic analysis

Phylogenetic relationships were reconstructed using a concatenated nucleotide dataset of 69 PCGs (64,654 bp) from 16 *Paeoniaceae* chloroplast genomes (Figure 6). The topologies of the ML and BI trees were highly congruent and strongly supported.

**Figure 6.**
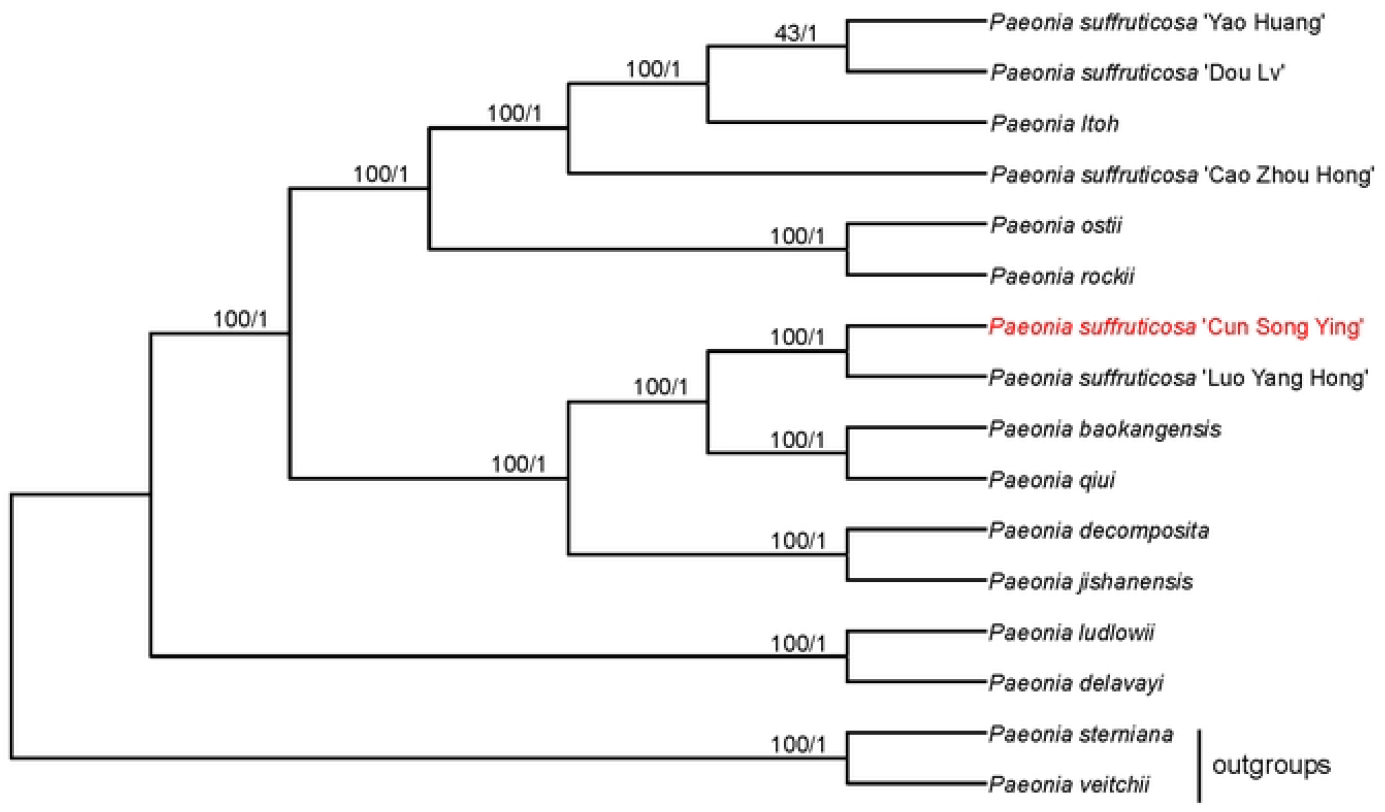
Phylogenetic tree of *Paeoniaceae* inferred from concatenated chloroplast CDS sequences of 69 PCGs. The tree was constructed using ML and BI methods. Numbers above branches represent bootstrap values and posterior probabilities.

Herbaceous peonies (*P. veitchii* and *P. sterniana*) formed distinct outgroups. All woody peonies clustered into a strongly supported monophyletic clade. Within this clade, *P. ludlowii* and *P. delavayi* diverged first, forming an independent lineage. The core woody peonies were divided into two major sister lineages.

The first lineage included wild ancestors such as *P. baokangensis, P. qiui, P. decomposita*, and *P. jishanensis*, together with the cultivated varieties *P. suffruticosa* ‘Luo Yang Hong’ and its closely related Japanese cultivar *Paeonia suffruticosa* ‘Cun Song Ying’. The second lineage comprised both wild and cultivated taxa, including *P. ostii* and *P. rockii*, along with well-known cultivars such as *P. suffruticosa* ‘Yao Huang’, ‘Dou Lv’, ‘Cao Zhou Hong’, and interspecific hybrids such as *Paeonia Itoh*.

This phylogenetic framework highlights the multiple hybrid origins of cultivated peonies and strongly supports the extremely close genetic relationship between *Paeonia suffruticosa* ‘Cun Song Ying’ and *Paeonia suffruticosa* ‘Luo Yang Hong’.

## 4 Discussion

This study presents the first complete chloroplast genome of *Paeonia suffruticosa* ‘Cun Song Ying’, including its sequencing, assembly, and annotation, followed by a systematic analysis of genome architecture, repeat content, codon usage bias, and phylogenetic placement. Our findings enrich chloroplast genomic resources for *Paeoniaceae* and provide new insights for cultivar identification, phylogenetic reconstruction, and plastid evolutionary mechanisms.

### 4.1 Structure and conservation of the ‘Cun Song Ying’ chloroplast genome

The chloroplast genome of ‘Cun Song Ying’ is 152,704 bp and displays the canonical circular quadripartite structure comprising the LSC, SSC, and two IR regions. We annotated 134 functional genes, including 87 protein-coding genes, 39 tRNA genes, and 8 rRNA genes. Gene content, order, and copy number closely match those reported for other *Paeonia* species [34], indicating strong overall conservation of chloroplast genomes within *Paeoniaceae*. This pattern is consistent with the structural stability of plastid genomes across core eudicots.

IR regions are widely considered critical for maintaining chloroplast genome stability. In most plants they are highly conserved. For example, Xiao et al. (2025) examined 21 *Camellia* species and reported conserved SC/IR boundaries across the genus [35]. Similarly stable IR regions have been documented in *Capsicum* [36], Sapindaceae [37], and wild *Prunus* species [38]. In our analysis, boundary genes at the IR junctions (such as *rps19, trnH, ycf1*, and *ndhF*) and their positions showed no notable variation among *Paeonia* taxa, supporting strong IR conservation among close relatives. By contrast, the LSC and SSC regions exhibited greater sequence variability, particularly in noncoding intervals, highlighting promising loci for developing high-resolution molecular markers.

### 4.2 Sequence variation and identification of hypervariable regions

Nucleotide polymorphism (π) analyses showed higher conservation in coding regions than in noncoding regions, in line with trends observed in most plant chloroplast genomes. π values in the IR were markedly lower than those in the LSC and SSC, reinforcing the stabilizing role of the IR.

We identified six hypervariable regions (π > 0.01), spanning coding and intergenic segments associated with *psbA, ycf4, psbJ–psbL–psbF–psbE, psaJ–rpl33–rps12, rpoA– rps11*, and *ndhF*. These regions also display elevated variability in related taxa and thus represent candidate DNA barcodes for cultivar identification and population genetic studies in peonies [39, 40]. Notably, *ycf1* and *ndhF* carry strong phylogenetic signals across diverse plant groups [41, 42], making them suitable for resolving interspecific relationships.

### 4.3 Repeat elements and SSR distribution

We detected 43 dispersed repeats in ‘Cun Song Ying’ comprising 21 palindromic and 22 forward repeats, with no reverse repeats identified. Repeat lengths clustered within 30–35 bp and 39–54 bp, mirroring patterns seen in other *Paeonia* species. Such repeats may facilitate homologous recombination and contribute to sequence variation, thereby influencing chloroplast genome evolution.

SSR analysis revealed 73 loci, dominated by mononucleotide repeats (49), followed by dinucleotide (12), trinucleotide (7), and tetranucleotide (5) repeats. The number of SSRs varies among *Paeonia* species (approximately 55–75), a polymorphism useful for cultivar discrimination and genetic diversity assessment. Pentanucleotide SSRs were rare and detected only in a few species, suggesting possible lineage specificity.

### 4.4 Codon usage bias and evolutionary implications

Codon usage analysis showed a pronounced preference for A/U-ending codons, a common pattern in land plants likely shaped by combined effects of mutational bias and natural selection. Leucine (Leu) was the most frequently encoded amino acid, whereas cysteine (Cys) was least frequent. The RSCU value for TTA (Leu) reached 1.97, indicating strong preferential use.

These biases may reflect optimization of translational efficiency and accuracy over long-term evolution. A/U-ending codons may better match the chloroplast tRNA pool and translation machinery, potentially enhancing expression efficiency. Understanding codon usage patterns helps elucidate molecular evolutionary dynamics and can inform strategies to optimize heterologous gene expression in chloroplasts.

### 4.5 Phylogenetic relationships and implications for cultivar origins

Phylogenies inferred from 69 protein-coding genes using ML and BI strongly support a sister relationship between *Paeonia suffruticosa* ‘Cun Song Ying’ and the cultivated *P. suffruticosa* ‘Luo Yang Hong’ (bootstrap = 100), indicating extremely close relatedness. This genome-scale evidence suggests tight genetic connections between the Japanese-introduced ‘Cun Song Ying’ and peony cultivar groups from central China, consistent with historical cross-regional introductions and exchange.

Woody peonies formed a well-supported monophyletic group that segregated into two major clades. Clade I included wild species such as *P. baokangensis, P. qiui, P. decomposita*, and *P. jishanensis*, together with cultivars including ‘Luo Yang Hong’ and ‘Cun Song Ying’. Clade II comprised *P. rockii, P. ostii*, and their derivative cultivars such as ‘Yao Huang’, ‘Dou Lv’, and Itoh hybrids, a lineage that appears to have evolved independently in both genetic and morphological traits relative to Clade I.

In summary, our chloroplast genome analyses clarify the close affinity between *Paeonia suffruticosa* ‘Cun Song Ying’ and ‘Luo Yang Hong’, providing a theoretical basis for peony cultivar identification and genetic provenance. Future integration with nuclear genomic data will enable deeper reconstruction of hybridization and domestication histories.

## 5 Conclusion

This study reports the first complete chloroplast genome of *Paeonia suffruticosa* ‘Cun Song Ying’, filling an important gap in genomic resources for this cultivar. The identified hypervariable regions and SSR markers offer valuable tools for peony cultivar identification, genetic diversity assessment, and conservation research. In addition, codon usage bias analysis provides a theoretical foundation for future chloroplast genetic engineering in peonies.

Future work should expand sampling to incorporate more accessions and apply population genomics approaches to elucidate the domestication history and dispersal patterns of peony cultivars. Integrating comparative transcriptomics and functional genomics will further clarify the roles of chloroplast genes in shaping traits such as flower color and stress resistance, thereby providing new molecular targets for breeding and genetic improvement.

## Acknowledgment

The authors are very grateful to professor Changyong Gao (Heze University) for providing materials on *Paeonia suffruticosa* ‘Cun Song Ying’.

## Authorship contribution statement

Wenzhen Cheng: Conceptualization, Writing-original draft, Funding acquisition. Conghao Hong: Data curation. Mingyu Li: Methodology, Software, Writing-review & editing.

## Supplementary data

Supplementary Data are available.

## Data Availability Statement

The sequence data generated in this study are available in GenBank of the National Center for Biotechnology Information (NCBI) under the access number PX252292.

---

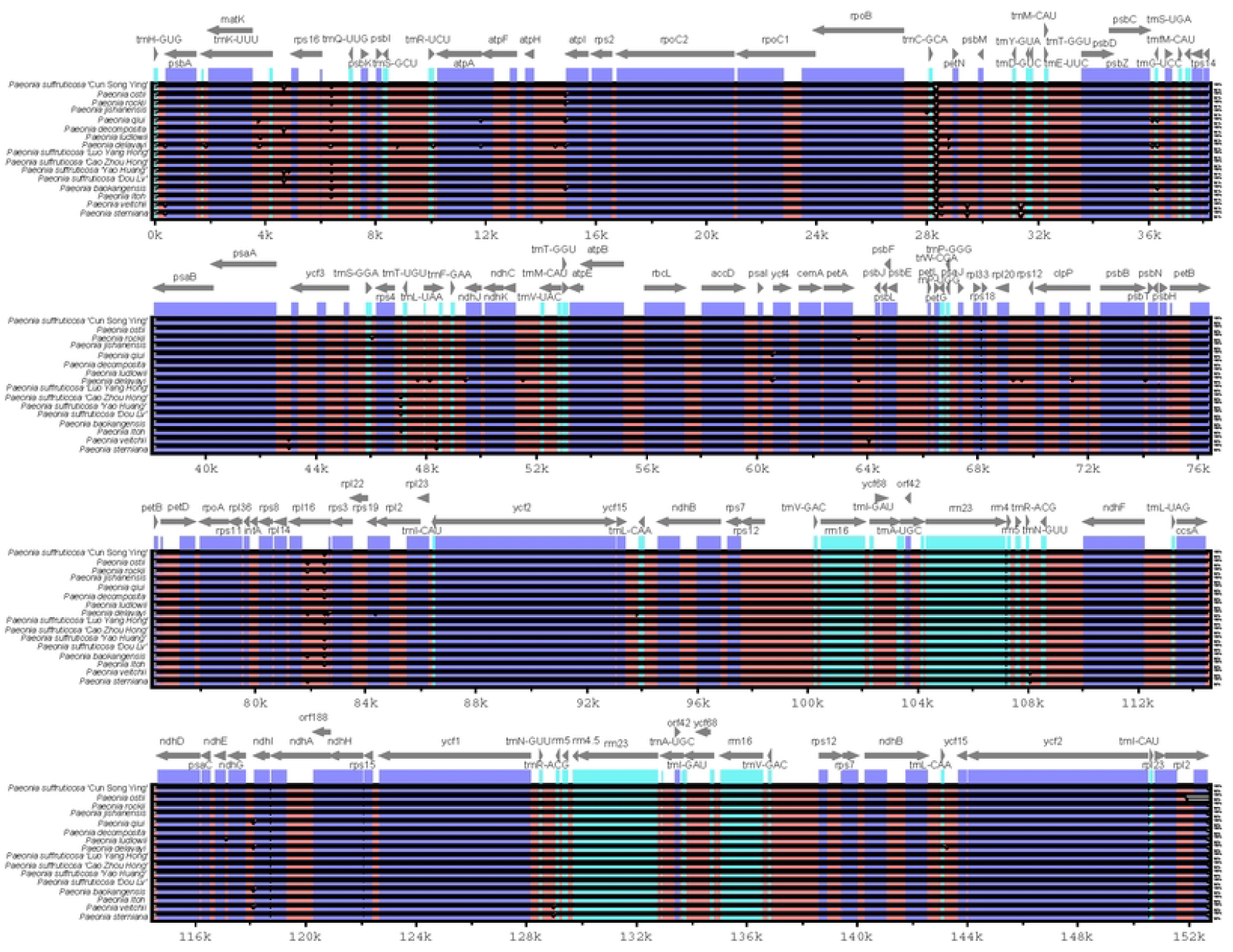

